# Minimal Data Needed for Valid & Accurate Image-Based fMRI Meta-Analysis

**DOI:** 10.1101/048249

**Authors:** Camille Maumet, Thomas E. Nichols

**Affiliations:** Warwick Manufacturing Group, The University of Warwick, Coventry, UK.; Statistics Department, The University of Warwick, Coventry, UK.

## Abstract

Meta-analysis is a powerful statistical tool to combine results from a set of studies. When image data is available for each study, a number of approaches have been proposed to perform such meta-analysis including combination of standardised statistics, just effect estimates or both effects estimates and their sampling variance. While the latter is the preferred approach in the statistical community, often only standardised estimates are shared, reducing the possible meta-analytic approaches. Given the growing interest in data sharing in the neuroimaging community there is a need to identify what is the minimal data to be shared in order to allow for future image-based meta-analysis. In this paper, we compare the validity and the accuracy of eight meta-analytic approaches on simulated and real data. In one-sample tests, combination of contrast estimates into a random-effects General Linear Model or non-parametric statistics provide a good approximation of the reference approach. If only standardised statistical estimates are shared, permutations of z-score is the preferred approach.

## 1 Introduction

A growing literature is focusing on the lack of statistical power in neuroimaging studies (see, e.g. [2]), feeding the debate on the validity and reproducibility of published neuroimaging results. Meta-analysis, by providing inference based on the results of previously conducted studies, provides an essential method to increase power and hence confidence in neuroimaging.

A number of methods have been proposed for neuroimaging meta-analysis (see [10] for a review). As the results of neuroimaging studies are usually conveyed by providing a table of peak coordinate and statistics, most of these metaanalyses are restricted to combining coordinate-based information. Nevertheless the best practice method is an Intensity-Based Meta-Analysis (IBMA) that combines the effect estimates and their standard errors from each study [1].

In order for IBMA to be possible in neuroimaging, tools for sharing 3D volumes obtained as a result of a statistical analysis are needed. Various efforts are currently underway to facilitate sharing of neuroimaging data but emphasis is usually on statistical maps (see, e.g. [2]). There are three evident approaches to sharing summary data from each study *i*:

1. the contrast estimates *Y*_*i*_ and contrast variance estimates 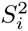.
2. the contrast estimates *Y*_*i*_.
3. the standardized statistical maps *Z*_*i*_.

Depending on how much data is shared, different strategies can be used to combine the available results into a meta-analysis. While the first option is the best practice, leading to statistically optimal estimates [4], working with contrasts requires fastidious tracking of data, model and contrast vector scaling. Given the growing interest in data sharing in the neuroimaging community, and the relative easiness of sharing just (unitless) statistic maps, there is a need to identify what is the minimal data to be shared in order to allow for future IBMA.

Here we compare the use of IMBA using 9 meta-analytic approaches: 2 approaches use *Y*_*i*_’S and 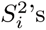, 2 *Y*_*i*_’S only and 5 *Z*_*i*_’S. We compare the validity and the accuracy of the eight meta-analytic approaches on simulated and real data including 21 studies of pain in control subjects.

Section 2 describes the meta-analytic estimates along with the experiments undertaken on simulated and real data to assert their validity. The results are described in section 3. Finally, we conclude in section 4.

## 2 Methods

### 2.1 Theory

For study *i* = 1,…, *k* we have contrast estimate *Y*_*i*_, its contrast variance estimate 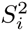 (i.e. squared standard error), its statistic map *Z*_*i*_, and its sample size *n*_*i*_.

#### Combining contrast estimates and their standard error

The gold standard approach is to fit contrast estimates and their standard error with a hierarchical general linear model (GLM) [4], creating a third-level (level 1: subject; level 2: study; level 3: meta-analysis). The general formulation for the study-level data is:

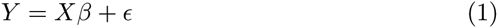

where *β* is the meta-analytic parameter to be estimated, *Y* = [*Y*_*i*_ … *Y*_*k*_]^*T*^ is the vector of contrast estimates, *X* is the *k* × *p* study-level matrix (typically just a column of ones) and *∈* ~ *N*(0, *W*) is the residual error term. Eq. (1) can be solved by weighted least square giving:

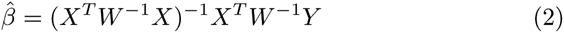

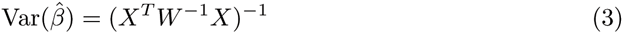

In a meta-analysis random-effects (RFX) model, we have 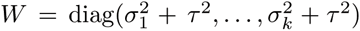 where *τ*^2^ denotes the between-study variance. Approximating 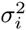 by 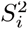 and given *τ̂*^2^ an estimate of *τ*^2^ we obtain the statistics detailed in Table 1 for one-sample tests. This reference approach will be referred to as **Mixed-effects (MFX) GLM**. In a **fixed-effects(FFX) GLM** (i.e. assuming no or negligible between-study variance), we have 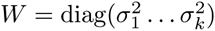 where 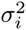 denotes the contrast variance for study *i*.

**Table 1.** Statistics for one-sample meta-analysis tests and their sampling distributions under the null hypothesis *H*_0_. Empirical null distributions are determined using permutations with sign flipping.

| | Meta-analysis statistic | Nominal $H_0$ distribution | Inputs |
| --- | --- | --- | --- |
| FFX GLM | $\left(\sum_{i=1}^k \frac{Y_i}{S_i^2}\right) / \sqrt{\sum_{i=1}^k 1/S_i^2}$ | $\mathcal{T}_{(\sum_{i=1}^k n_i - 1) - 1}$ | $Y_i, S_i^2$ |
| MFX GLM | $\left(\sum_{i=1}^k \frac{Y_i}{S_i^2 + \hat{\tau}^2}\right) / \sqrt{\sum_{i=1}^k 1/(S_i^2 + \hat{\tau}^2)}$ | $\mathcal{T}_{k-1}$ | $Y_i, S_i^2$ |
| RFX GLM | $\left(\sum_{i=1}^k \frac{Y_i}{\sqrt{k}}\right) / \hat{\sigma}_C^2$ | $\mathcal{T}_{k-1}$ | $Y_i$ |
| Contrast Permutation | $\left(\sum_{i=1}^k \frac{Y_i}{\sqrt{k}}\right) / \hat{\sigma}_C^2$ | Empirical | $Y_i$ |
| Fisher's | $-2 \sum_{i=1}^k \ln(\Phi(-Z_i))$ | $\chi_{(2k)}^2$ | $Z_i$ |
| Stouffer's | $\left(\sum_{i=1}^k Z_i\right) / \sqrt{k}$ | $\mathcal{N}(0, 1)$ | $Z_i$ |
| Weighted Stouffer's | $\left(\sum_{i=1}^k \sqrt{n_i} Z_i\right) / \sqrt{\sum_{i=1}^k n_i}$ | $\mathcal{N}(0, 1)$ | $Z_i, n_i$ |
| Z MFX | $\left(\sum_{i=1}^k Z_i\right) / \sqrt{k} \hat{\sigma}$ | $\mathcal{T}_{k-1}$ | $Z_i$ |
| Z Permutation | $\left(\sum_{i=1}^k Z_i\right) / \sqrt{k}$ | Empirical | $Z_i$ |

#### Combining contrast estimates

If the 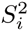 are unavailable, the contrast estimates *Y*_*i*_ can be combined by assuming that the within-study variance 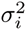 is roughly constant 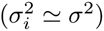 or negligible in comparison to the between-study variance 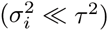. Then 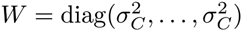 where 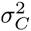 is the combined within and between subject variance, i.e. 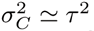 or 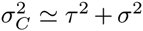 (note, however, in this setting we do not separately estimate *τ*^2^ or *σ*^2^). Under these assumptions, Eq. (1) can be solved by ordinary least squares giving:

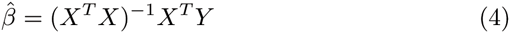

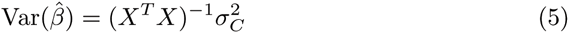

Given 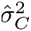 an estimate of 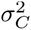 we obtain the statistics presented in Table 1 for one sample tests. This approach will be referred to **RFX GLM** in the following.

As an alternative to parametric approaches, non-parametric inference [6,9] can be performed by comparing the RFX GLM T-statistic to the distribution obtained “sign flipping”, i.e. randomly multiplying each study’s data by 1 or ‐1, justified by an assumption of independent studies and symmetrically distributed random error. This approach will be referred to as **Contrast permutation**.

#### Combining standardised statistics

When only test statistic images are available there are a several alternate approaches available. **Fisher’s** meta-analysis provide a statistic to combine the associated p-values [5]. **Stouffer’s** approach combines directly the standardised statistic [13]. In [14] following [7], the author proposed a weighted method that weights each study’s *Z*_*i*_ by the square root of its sample size [3,7]. This approach will be referred to as **Weighted Stouffer’s**. All these meta-analytic statistics assumes no or negligible between-study variance and are suited only for one-sample tests. The corresponding statistics are presented in Table 1. As suggested in [1], to get a kind of MFX with Stouffer’s approach, the standardised statistical estimates *Z*_*i*_ can be combined in an OLS analysis. The corresponding estimate, referred as **Z MFX** is also provided in 1

With contrasts, non-parametric inference [6, 9] can be obtained by sign flipping on the *Z*_*i*_’S. This approach will be referred to as **Z permutation**.

*Approximations* In practice, all of the methods based on contrast data have approximate parametric null distributions. The nominal distributions listed in Table 1 are under the (unrealistic) assumption of homogeneous standard errors over studies; even if all studies are ‘clean’ and conducted at the same center, variation in sample size will induce differences in 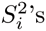. Further, even under homoscedasticity, MFX GLM’s null is approximate due to iterative estimation of *τ̂*^2^.

### 2.2 Experiments

#### Simulations

Due to the approximate nature of the sampling distributions, we conduct simulations to evaluate the validity of each estimator under inhomogeneity of contrast variances 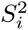 and under the presence of non-negligible between-study variance.

To verify the validity of each estimator under the null hypothesis we estimated the false positive rate at *p* < 0.05 uncorrected. For each meta-analysis, we simulated *Y*_*i*_ and 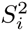 such as:

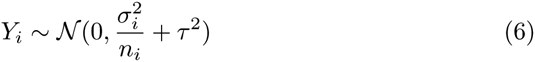

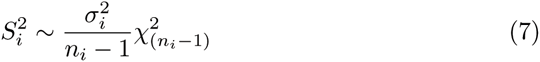

where of 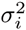 ∈ [1/2,1,2,4] is the within-study variance, *τ*^2^ ∈ [0,1/20] is the between-study variance (fixed-effects models are strictly only appropriate for *τ*^2^ =0). For different number of studies per meta-analysis we used: *k* ∈ [5,10, 25, 50], and set the number of subjects per studies *n*_*i*_ to vary across the common range of sample sizes in neuroimaging studies. In each simulated meta-analysis we simulated one study with exactly 20, 25, 10 and 50 subjects. For the remaining studies 1/4 of the *n*_*i*_’S were drawn from *U*(11,20), 1/4 from *U*(26,50) and the remaining from *U*(21,25), where *U*(a,b) is the discrete uniform distribution on the integers *a* to *b* inclusive. A total of 32 parameter sets 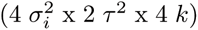 was therefore tested and a total of 71^3^ realisations were created.

#### Real data

We then compared the 8 meta-analytic estimators to the reference approach, MFX GLM, on a dataset of 21 studies of pain. Comparability of contrast estimates depends on equivalent scaling of the data, models, and contrast vectors. Data scaling was consistently performed by FSL, setting median brain intensity to 10,000; model were all created by FSL’s Feat tool; and contrasts were constructed to preserve units, with sum of positive elements equal to 1, sum of negative elements equal to ‐1.

To investigate the presence of between-study variation, we computed the ratio of the between-study variance (estimated using FSL’s FLAME [12]) to the total variance (sum of between‐ and within-study variances), as suggested in [3]. Here we use the average (across study) within-study variance as an estimate of within-study variance in the denominator: 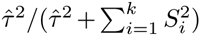 Using this metric, voxels with values close to 0 present negligible between-study variance and values close to 1 outline appreciable study heterogeneity and the importance of RFX models.

Then for each estimator we compared the standardised meta-analytic statistic to the z-statistic obtained with the reference approach. Overestimation of z-statistic leads to overly optimistic detections while underestimation outline a reduced sensitivity of the approach.

## 3 Results

### 3.1 Simulations

Fig. 1 displays the false positive rate at *p* < 0.05 obtained for the eight estimators over all set of parameters in the absence and presence of between-study variation. As expected, the fixed-effects meta-analytic summary statistics, i.e. Fisher’s, Stouffer’s and weighted Stouffer’s estimates, are liberal in the presence of study heterogeneity. The original Fisher’s approach is the most invalid. More surprising, FFX GLM is also invalid with homogeneous studies. The explanation is over-estimation of degrees-of-freedom (DF); while DF is computed as (∑*n* – 1) – 1, under heteroscasdicity (from *σ*_*i*_ or *n*_*i*_) it will be much lower [11]. Z MFX and GLM RFX provide valid estimates, and the permutation estimates are valid but tend to be conservative with greater variation in false positive rates.

**Fig. 1.**
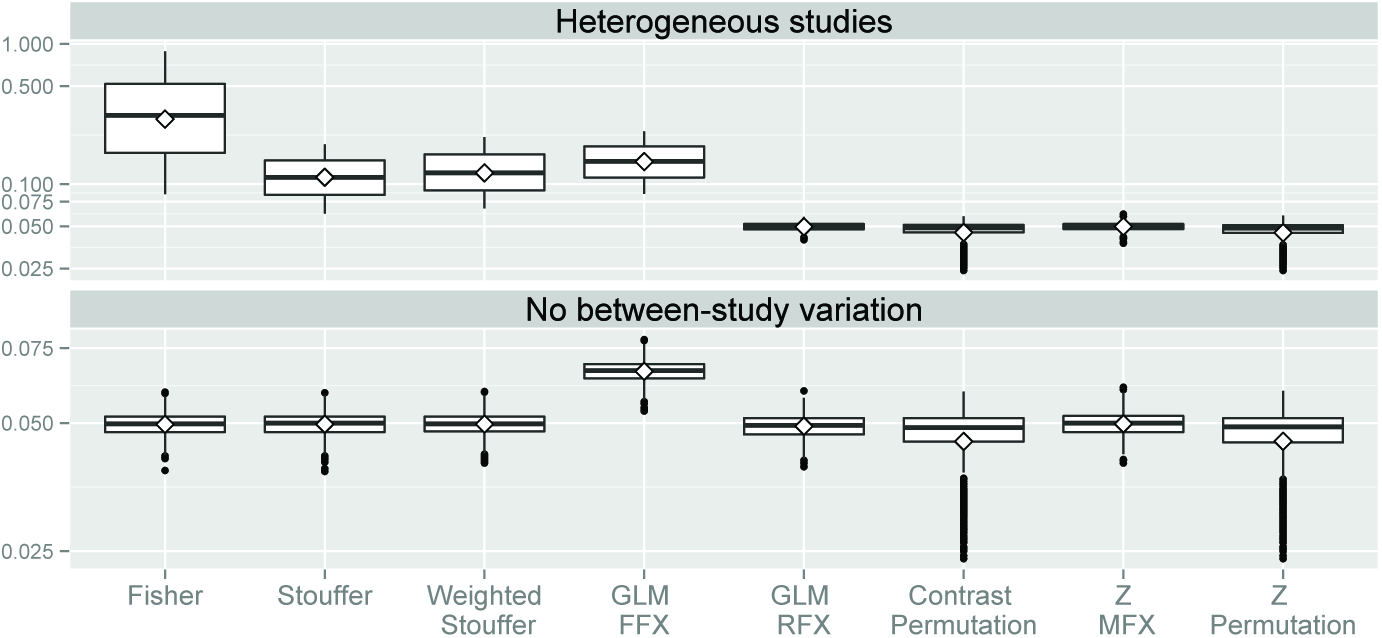
False positive rates of the meta-analytic estimators under the null hypothesis for *p* < 0.05.

The impact of the number of studies involved in the meta-analysis and of the size of the within-study variance are investigated in Fig. 2. Permutation inference is valid but conservative when 5 studies are used; this is because there are only 2^5^ = 32 possible permutations and thus 1/32 = 0.03125 is largest attainable valid P-value. All approaches perform equally as soon as 10 or more studies are included in the meta-analysis.

**Fig. 2.**
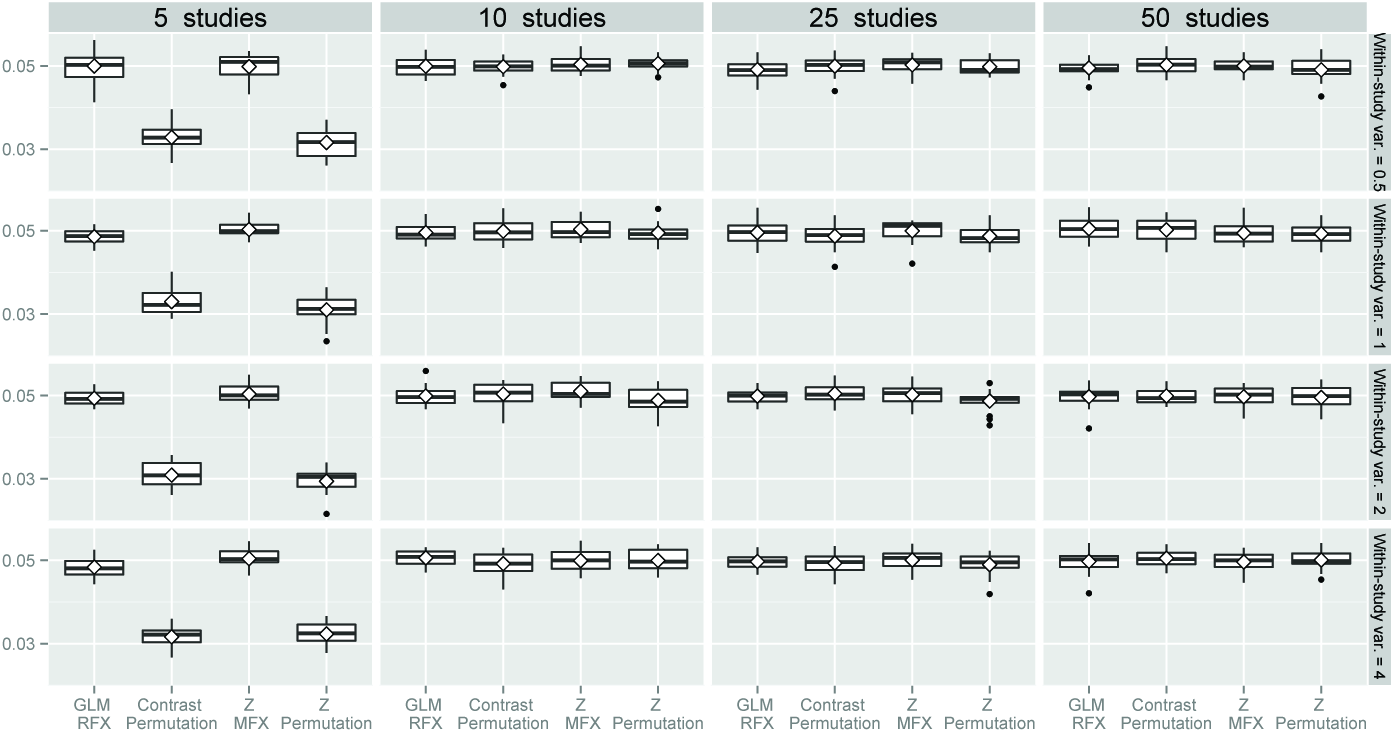
False positive rates of the RFX meta-analytic estimators under *Ho* for *p* < 0.05 as a function of the number of studies and the within-study variance.

### 3.2 Real data

The histogram of the ratio of between-subject variance to total variance is displayed in Fig. 3. From this graph it is clear that for most of the voxels the estimated between-study variance is greater than the within-study variance. We can therefore suppose the presence of study heterogeneity (non negligible between-study variance) in this collection of studies.

**Fig. 3.**
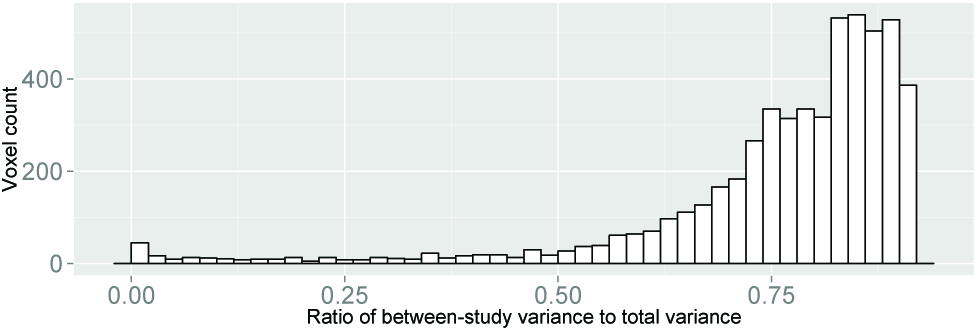
Histogram of the between-study variance to the sum of the between-subject variance and the mean within-study variance.

Fig. 4 plots the difference between the z-score estimated by each metaanalytic approach against the reference z-score computed with MFX GLM. All FFX statistics provide overly optimistic z-estimate suggesting, again, that study heterogeneity is present in the studied dataset. Among the RFX meta-analytic approaches, GLM RFX and contrast permutations provide z-scores estimate that are equal or smaller than the reference. Z permutation provides slightly larger z-scores between 1 and 3 (reference p-values between 0.16 and 0.0013) but is mostly in agreement with the reference z-scores. On the other hand, Z MFX is more liberal than the reference for z-score ranging from 3 to 5 (reference p-values between 0.0013 and 2.9e-07) and more stringent for z-scores smaller than 5.

**Fig. 4.**
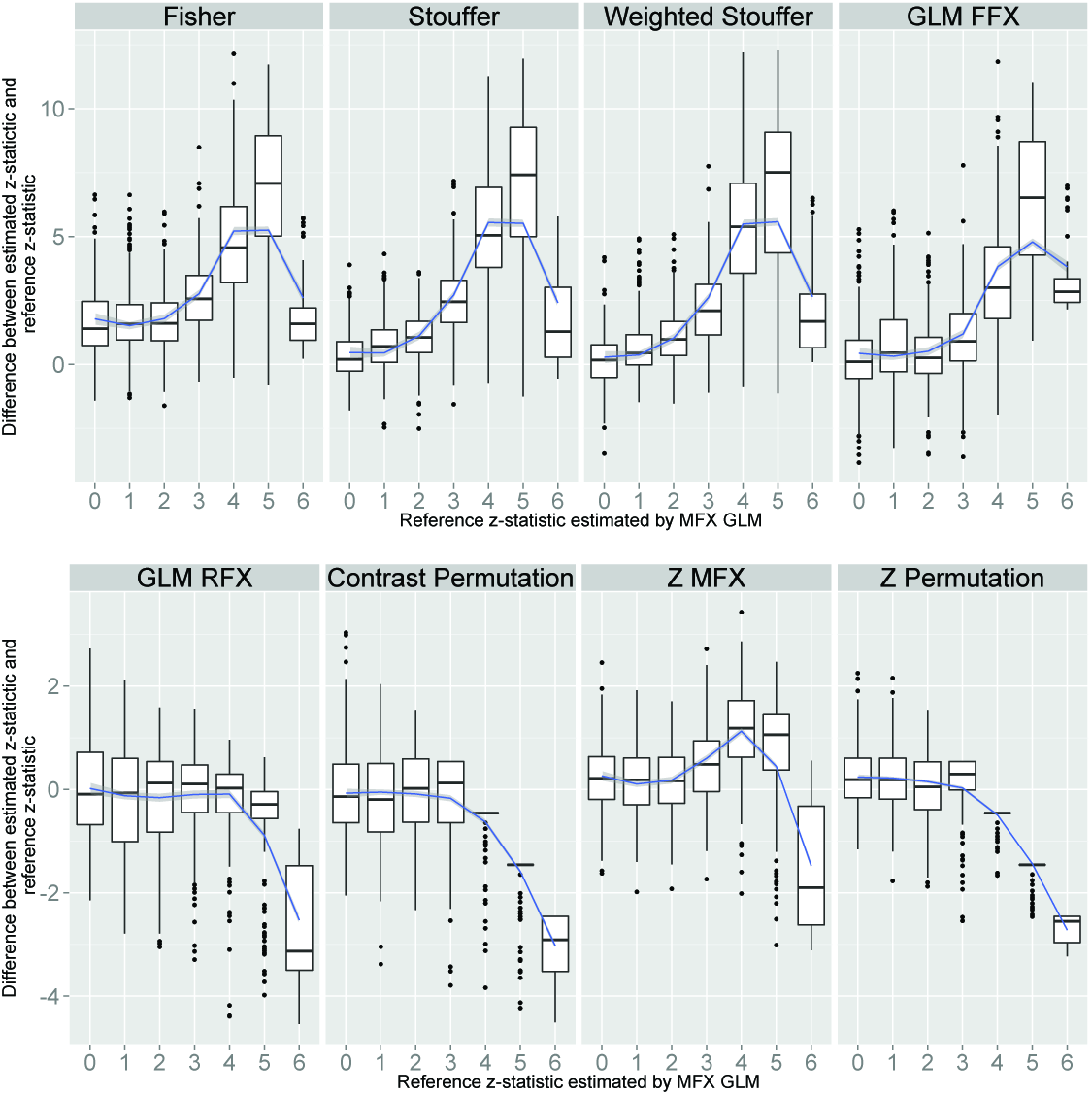
Difference between the z-score estimated from each meta-analytic approach and the reference z-score from MFX GLM as a function of reference z-score.

## 4 Conclusion

We have compared eight meta-analytic approaches in the context of one-sample test. Through simulations, we found the expected invalidity of standard FFX approaches in the presence of study heterogeneity, but also of FFX GLM even with no between-study variation. In a real dataset of 21 studies of pain, there was evidence for substantial between-study variation that supports the use of RFX meta-analytic statistics. When only contrast estimates are available, RFX GLM was valid. This is in line with previous results on within-group one-sample t-tests studies [8]. When only standardised estimates are available, permutation is the preferred option as the one providing the most faithful results. Further investigations are needed in order to assess the behaviour of these estimators in other configurations, including meta-analyses focusing on between-study differences.

## 5 Acknowledgements

We gratefully acknowledge the use of this data from the Tracey pain group, FMRIB, Oxford. CM and TEN were supported by the Wellcome Trust.

